# Social copying drives spatial allocation in gregarious herbivores

**DOI:** 10.64898/2026.11.03.764068

**Authors:** Jan Lewenstein-Sanpera, Daniel Campos, Joan Bauzà, Daniel Oro

## Abstract

Resting patch selection in free-ranging social cattle (*Bos taurus*) inhabiting steep, rugged, mountain valleys is critical, because the recurrent use of flatter resting patches may reduce energetic costs and enhance fitness prospects. As these patches may constitute a limited resource, understanding their pattern of use dynamics may help explain resting patch selection. We therefore combine empirical data with a theoretical modelling framework to investigate the processes underlying resting patch selection. Using high-resolution GPS tracking data from 144 cows collected over three seasons, we identify several recurrently used resting patches located on significantly gentler slopes than expected in the valley. We then build a simple, memory-free agent-based model, which suggests that the dynamics of resting patch selection arise from social copying and, to a lesser extent, from individual heterogeneity. The model quantitatively reproduces three empirical observables: a slowing (sub-linear) discovery of new sites through time, heterogeneous occupancy across resting patches and similarity in patch use among individuals. These results suggest that collective dynamics in social species can emerge primarily from a simple social mechanism with weaker modulation from inter-individual heterogeneity, such as age, without explicitly incorporating individual memory to reproduce the aggregate patterns.

## I. INTRODUCTION

Animals face complex decision-making challenges, where the choices on where to move and how to allocate time and energy to acquire resources have direct consequences for survival and fitness [1]. Movement, as a consequence of the spatial and temporal dynamic heterogeneity of the resource [2], reproduces a glimpse of these situations, integrating both environmental signals and internal states (such as a cliff in the way, a predatory scent ahead or a reproductive drive). Consequently, it is important to understand the patterns that arise during animal movement and interpret their ecological and evolutionary significance [1].

In social species, this decision-making is strongly shaped by social interactions and relies on two main contributions [3–5]: on the one hand, there is the public information due to the presence of conspecifics in the same resource patch, and it gives rise to behaviours such as collective responses. This information is available to the entire social group as a unit, while, on the other hand, there is the private information, which is gathered independently by each individual [6]. Both factors produce a feedback loop where individual behaviour scales up to determine social structures, while at the same time these structures drive individual decisions [4, 7]. The resulting dynamics are therefore greatly influenced by collective decisions and not just private information [8, 9]. For example, behaviours such as social copying, where individuals copy a widely distributed behaviour and are density-dependent, have been observed to play a significant role in decision-making, especially under environmental perturbations [10, 11]. This flexibility in behaviour and feedback between private and public information often optimises fitness [11] and may help explain the persistence of such behaviours from an evolutionary perspective [12, 13].

Private information should as well include the spatiotemporal dynamics of resources, such as the spatial configuration of the habitat, allowing individuals to respond to environmental stochasticity [14, 15]. This, in turn, will shape social interactions and structure [4, 16, 17] and potentially provoke different dynamical patterns linked to specific resources [18, 19]. On this matter, the simplest case relies on fixed (both in time and space) resources, such as resting patches where animals can rest safely, reduce energetic costs, and minimize mortality risk . These offer the possibility to study the observed dynamical patterns in a controlled environmental setup, where spatial constraints are only given by the topography, rather than by their stochastic nature.

In this context, animals integrating the spatiotemporal dynamics can show signs of memory-driven responses [20, 21]. Such behaviours are characterised by persistent patterns, far away from stochastic responses [1], with higher spatio-temporal correlations [22] and better predictability in the statistical sense. For example, one can consider preferential returns as the spatial fidelity to a particular patch, which can be studied as the accumulation of revisits to specific sites with a deviation from a linear occupancy distribution [23– This trait, usually associated to memory, involves an energy trade-off between exploring new sites (exploration) and revisiting known patches (exploitation) [27]. However, as mentioned above, decision-making is tightly correlated to the social structure the group possesses [4, 13], and there are examples of socially triggered, instead of memory, preferential returns[10, 28].

Here, we explore how a large social herbivore exhibits a preferential return trait linked to the resting patch selection and we assess whether this behaviour is modulated by individual heterogeneities such as age and degree of social interaction. We hypothesise that the resting patches are specifically selected to be topographically flatter and propose that this observed behaviour can emerge from a simple social copying mechanism, due to the nature of the herbivore’s interactions in the valley, instead of individual memory. Moreover, we believe that the heterogeneities between individuals encode the necessary information for a model to reproduce the observed behavioural patterns.

This article is structured as follows: In section II: Methods we describe the study data set and statistical processing, it is followed by section III: Theoretical Model which describes the used model along with its main characteristics. Then section IV: Results includes both the empirical and the model’s performance and is followed by section V: Discussion. Finally, there is a brief section VI: Conclusions to close the arguments.

## II. METHODS

### A. Biological model and procedures

We studied a population of free-ranging cattle (*Bos taurus*), which are highly social animals that retain their “wild” behaviour even if domesticated and exhibit developed memory traits [24, 29]. This population lives in a mountain valley in the Catalan Pyrenees, the Catllar Valley (*42°22’55*.*34”N, 2°15’19*.*77”E*), which spans an area ≈ 13 *km*^2^ and varies in altitude from ca. 1400 metres, where an electric fence prevents cattle from going lower, to ca. 2700 metres. The herd enters the study valley every year at the beginning of July and remains there until late October, when they are led to lower elevations. Individuals do not change year after year, except due to mortality causes, and are born during the summer period in the study valley. Our own observations show that losing balance, with the potential of falling and rolling down the slopes, is the main cause of mortality in the herd. For this reason, we focus our study on the critical resource of the resting patch, which allows cattle to spare energy and reduce mortality, thus playing a critical role in decision-making with fitness consequences.

The datasets used for this study consist of GPS cattle positions for the free-ranging period (from mid June to mid October) from 2021 to 2023. Each data set is then analysed to extract the night resting positions, based on GPS accelerometer and daily data, to correctly identify cattle behaviour [30]. After cleaning out those data points consisting of malfunctioning GPS or cattle that mainly range outside of the study area, a total amount of 144 out of the initial 245 cow series are considered throughout the study period, some of these for the same individuals across the multiple years of the study. Data included positions (latitude and longitude) spaced every 10 *min* for the whole study period coupled with raster inferred altitude, giving a real estimate of the individual position in time and space.

## III. THEORETICAL MODEL

In order to interpret the statistical patterns associated to the movement between resting patches, we introduce a minimal model which incorporates imitation effects in the dynamics of decision making each individual cow makes every night. In biological terms, this corresponds to individuals either copying the resting choices of nearby conspecifics or exploring alternative patches independently. For this, we will use the noisy Voter Model [31] (also known as the Kirman’s model in the context of economics [32, 33]), which represents a standard mathematical reference in the context of social imitation. In this model, one has a group of *N* individuals which have to choose sequentially between *M* different options (e.g patches). To take their decisions, the individuals can either imitate other individuals or take their decisions at random. In the present work, the choice mechanism is assumed to follow these rules:

for *i* from 1 to *N* do:

1. With probability *p*_*i*_ the individual *i* chooses the resting site currently used by another individual *j* (*imitation rule*), with *j* chosen at random between a set Ω_*i*_ of neighbour individuals.
2. Otherwise (with probability 1 ™ *p*_*i*_) the individual *i* chooses among the *M* available patches at random with uniform probability (*exploration rule*), repre-senting uninformed exploration.

So that, we apply sequentially the algorithm to every individual, so a whole time step (corresponding to one night in the context of cows) is completed once every individual has taken a decision.

Compared to the most standard version of the noisy Voter Model, our algorithm introduces two relevant generalisations. First, instead of assuming that any option is accessible to the individuals through imitation, we introduce the restriction that individuals to imitate must lie within a neighbourhood Ω_*i*_ of the current site where the individual is. So that, prospection and imitation are assumed to be rather local. Specifically, we assign a label *k* = 1, 2, … , *M* to every site, so we assume that the neighbourhood Ω is defined by the condition |*k*_*i*_™*k*_*j*_| *< d*, where *k*_*i*_ and *k*_*j*_ are the labels for the current site of the individuals *i* and *j*, respectively. So, imitation is only possible if the sites of the two individuals are within a topological distance *d*. This reflects the assumption that imitation is spatially local, as individuals are more likely to copy nearby conspecifics than distant ones.

Second, we explicitly consider individual variability through the imitation probability *p*_*i*_. So that, individuals with lower *p*_*i*_ have a lower tendency to imitate (e.g. more experienced individuals), whereas those with higher *p*_*i*_ are more likely to rely on social information. In our case, we will take the *p*_*i*_’s to be Gaussian variables distributed around a mean value *p* with a standard deviation *σ*_*p*_ (values of *p*_*i*_ smaller than 0 or higher than 1, that make no sense, are then reassigned values *p*_*i*_ = 0 and *p*_*i*_ = 1, respectively). Importantly, to test whether social interactions and individual heterogeneity can explain the observed patterns, the model does not include explicit individual memory, so repeated use of patches would arise from social interactions, heterogeneity, and stochastic exploration.

### A. General dynamics of the noisy voter model

For the sake of clarity, let us review the most significant results known for the basic version noisy Voter Model where all individuals have identical behaviour (it is, *p*_*i*_ = *p* for any individual) and imitation is not restricted to local neighbours. We here just present an intuitive and schematic overview of the model, while a more formal and detailed description can be found elsewhere (see, e.g. [34–36]).

If we define *f* (*m, t*) as the average number of individuals at site *m* at time *t*, we can write

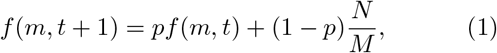

so in the stationary state where *f* (*m, t* + 1) = *f* (*m, t*) we obtain

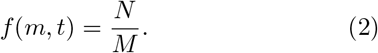

Accordingly, there are no preferential sites in the model on average. The main interest of the model, however, lies in the distribution of occupied sites that is reached in that stationary regime. When the probability of random choices is below a certain critical threshold *p*^*\**^ (which satisfies *p*^*\**^ *>* 1 ™ 1*/N* , then the dynamics is dominated by imitation and so consensus, or alternatively partial consensus (this is, a frozen state with all individuals sharing one common option, or maybe a small set of them) is observed in the stationary regime. Instead, for *p < p*^*\**^ random choices dominate the system dynamics, so the system is not characterized by a permanent or frozen state. Instead, individuals continuously jump from one option to another and so there are no sites with a much higher occupation than others.

To illustrate this idea formally, we introduce the occupation function *c*_*m*_(*n, t*), this is, the probability that *n* individuals choose site *m* at time *t*. This magnitude satisfies a master equation of the type

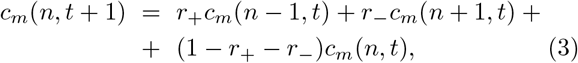

with

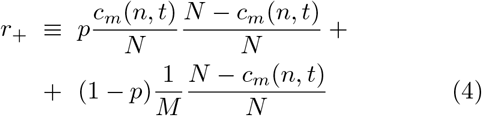

and

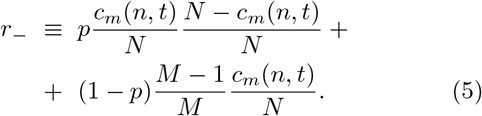

In expression (Equation 3), the first term on the right side corresponds to the case where an individual which was not previously at site *m* switches to it. The second term corresponds to the probability that the individual, which was previously at *m*, chooses this time a different resting site. Finally, the last term corresponds to the case where the individual does not change its site and so the occupation function remains the same.

Expanding the Master equation above for the case of large times and a large number of individuals (this is, *t* ≫ 1 and *N* ≫ 1, which corresponds to the so-called Kramers-Moyal expansion in statistical physics) yields

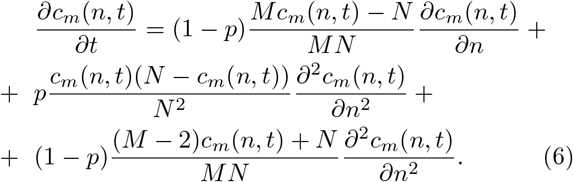

To clarify understanding of this equation, let us note that the expression Equation 6 can be rewritten as

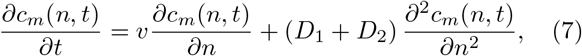

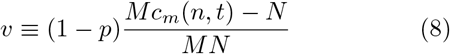

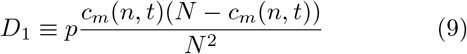

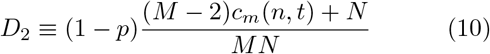

From these expressions, it can be seen that the drift coefficient *v* essentially drives the system towards *c*(*m, t*) = *N/M* (this is, towards a homogeneous state) due to exploration rule that appears with probability (1 ™ *p*). Instead, *D*_1_ and *D*_2_ correspond to fluctuating forces due to imitation and exploration, respectively. The first one vanishes only when either *c*(*m, t*) = 0 or *c*(*m, t*) = *N* , so it tries to lead the system towards the states where either the site is empty, or occupation is maximum (so promoting *consensus*). The second one is just an additional fluctuation that prevents the system from reaching absolute consensus, so even when *D*_1_ vanishes some level of fluctuations is always present in the dynamics.

Finally, we can ask what is the probability that an individual has visited *s* different patches at time *t*. While this is difficult to predict for the case with exploration, it becomes relatively easy for the case *p* = 1. Note that for this case only a subset (of size *M*_0_ ≤ *M*) of the patches is available at practice to the individuals (given by the set of patches occupied at *t* = 0). The probability that a new site is visited by the individual for the first time at time step *t* corresponds to the imitation probability *p* times the probability of choosing one among the *M*_0_ patches that has not been visited yet. Using combinatorics, the probability that after *t* nights the number of visited patches is *s* reads

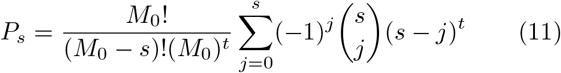

and the average value of patches visited is then

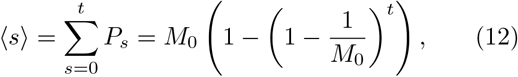

which grows from ⟨*s*⟩ = 0 (for *t* = 0), up to ⟨*s*⟩ → *M*_0_ (as *t* → ∞), as expected.

## IV. RESULTS

### A. Empirical results

We start by analysing the spatial resting locations for *N* = 144 individual data series, and obtain that these are specifically selected showing a different topographical feature than the rest of the valley. In Figure 1 we observe a comparison between the slope distribution of resting locations against a single random sampling throughout the study area. A Monte Carlo sampling method (with 1000 subsets) was applied to test the median slope distribution of the null models against the median of the resting locations, resulting in a statistically flatter slope for the resting locations (*p-value <* 0.001) and indicating that cattle tend to select flatter areas.

**FIG 1.**
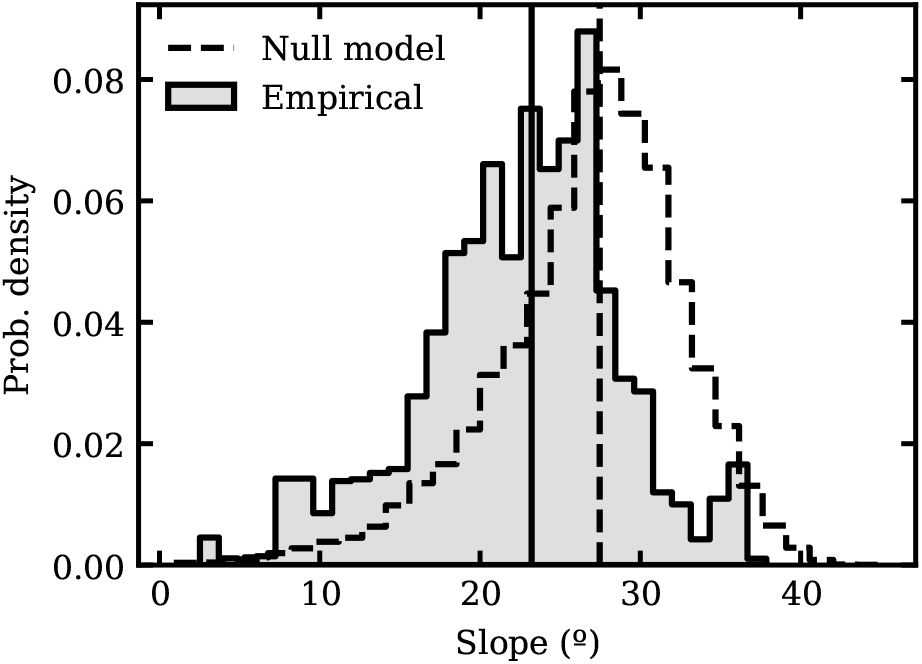
Observed resting patches slope distribution for the cattle population with respect to a null model considering random locations in the valley (we assume that cattle can explore the whole available space) in the study area. Medians are marked with vertical lines: empirical data median = 23.22 while null model data median = 27.47. Through a Monte Carlo sampling method, the null models confidence interval at (95%) is given by [27.37, 27.56] with the empirical median falling outside.

For every night in the study period, a single resting location per individual cow was selected. This was based on identifying a stationary location during the night considering the resting behaviour criterion introduced in [30]. Further analysis shows that cattle tend to cluster in fragmented recurrent patches Figure 2, which, through field observations, we can assume to have no maximum capacity and be reachable to all individuals. These patches are what we call the resting patch resource and, together with the topographical information, we infer that these sites are specifically selected. By studying the spatio-temporal dynamics associated to this resource (e.g. the movement between patches per night), the data suggest that the visiting rates to different patches (VUP) follows a sub-linear growth Figure 3 with cows exploring new sites with a time dependence as *t*^*α*^ where *α* = 0.619 ± 0.005. This sub-linear growth indicates a deviation from the stochastic response, consistent with a progressive shift from exploration to repeated use of known patches. Moreover, we also observe that patch saturation has not yet occurred, since the curves do not completely flatten out at *M* = 33, but its effect can be observed with a reduction of the slope (less available patches as time grows).

**FIG 2.**
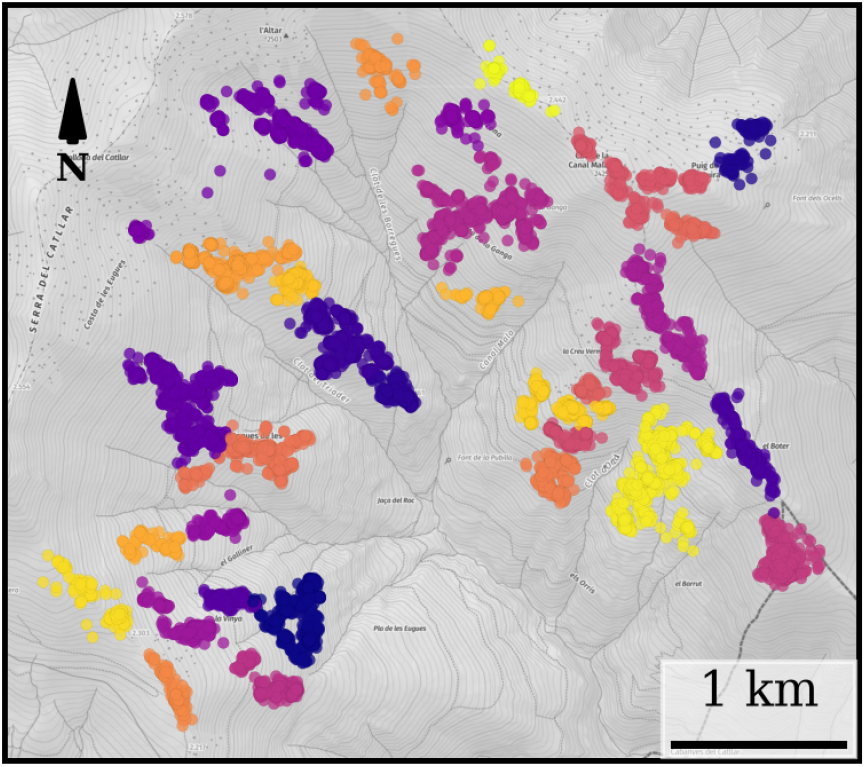
Observed resting patches computed with a clustering algorithm based on hierarchical density differences (HDB-SCAN). We assumed that individuals occupied a single patch per night and imposed that the patches have at least 80 points and these points are separated at furthest 90 meters to be considered in the same cluster. This resulted in 33 resting patches and 6.87% of outliers, with each colour representing a different patch while each point is a recorded night for a specific individual. Note that this is a 2D representation of the valley and clusters might appear closer than they are, with boundary points pertaining to the most probable cluster [37]. Outliers are omitted. Basemap provided by [38].

**FIG 3.**
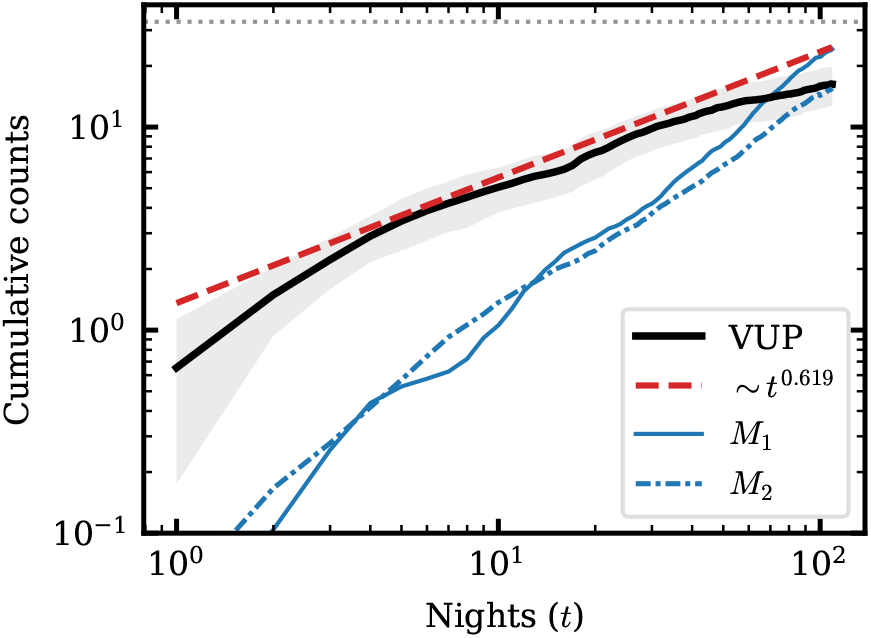
Cumulative count of the unique resting patches that were visited per cow per night (VUP) with their standard deviation cloud in gray. This VUP curve was fitted with a power law, resulting in *V UP* ∼ *t*^0.619^. In blue are represented the curves corresponding to *M*_1_ and *M*_2_; which represent the average occupancy over time of the most visited and second-most visited resting patches per cow respectively (standard deviation was omitted for the simplicity of the plot). The gray dotted line represents the total amount of patches, *M* = 33 in our system, as in Figure 2. Both axes in logarithmic scale.

Another interesting aspect, is that individual cows select specific patches with a higher probability than the rest. This can be seen with the evolution for *M*_1_ and *M*_2_, which represent the most and second-most visited resting patch per individual, respectively (the specific site changes between individuals). This is depicted in Figure 3. By the end of the time period, *M*_1_ = 24 ± 8, while *M*_2_ = 16 ± 4, clearly larger than the expected value if no site was preferred: *M*_1_ ≈ … ≈ *M*_*M*_ ≈ 1*/*33 * *nights* ≈ 5, which shows how preferential returns emerge in each in-dividual, although highly variable.

In order to understand the reasons behind this nonstochastic exploration behaviour and the preferential returns, we explored how individual heterogeneities, such as the cow’s age and sociality (an index computed as the average amount of individuals each cow spends the night with) affect the resting patch usage Figure 4. Our data suggest that these factors are statistically significant but play a minor role in determining the observed movement patterns.

**FIG 4.**
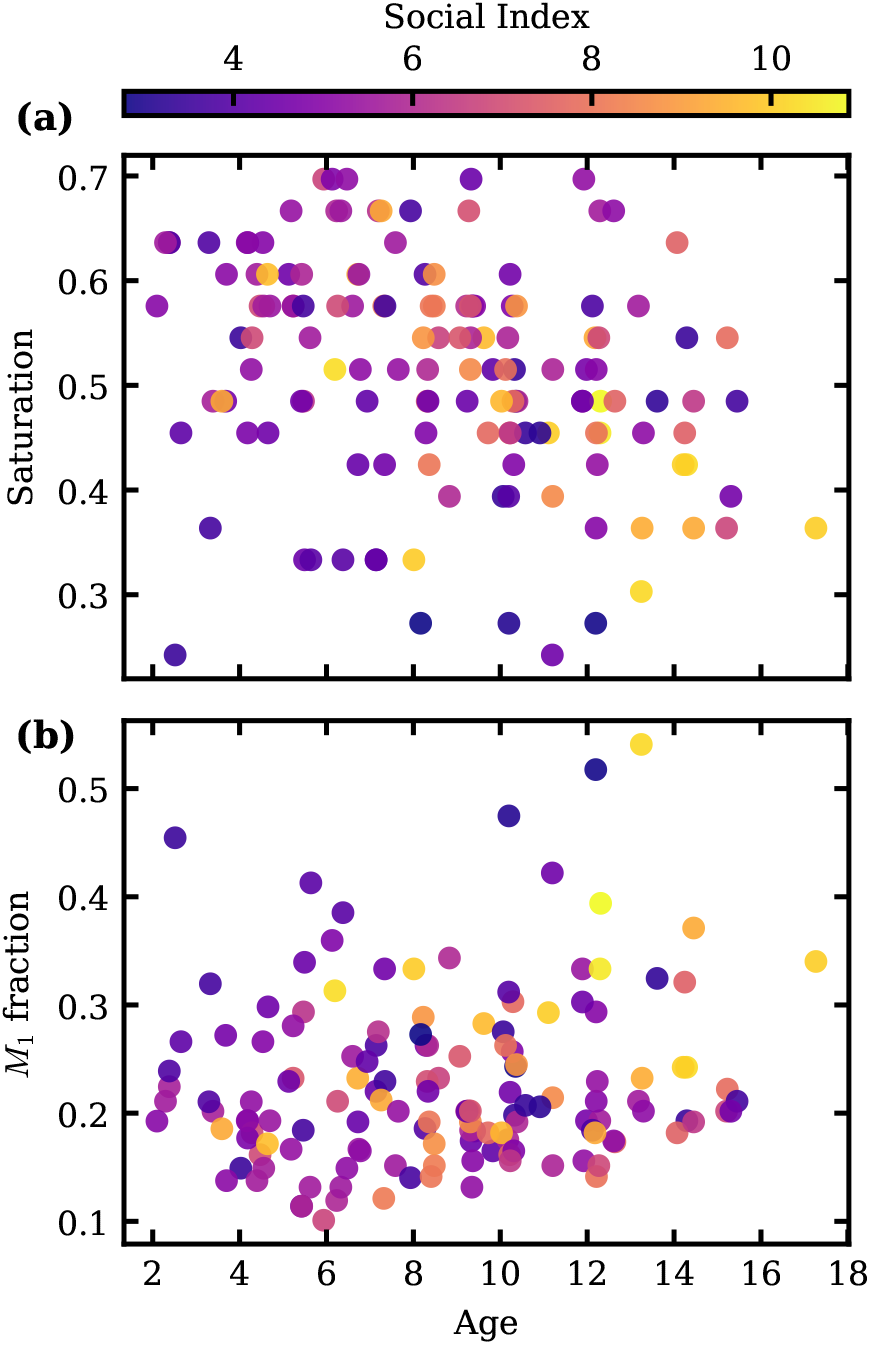
We show the dependency with age and sociality of the saturation, understood as the amount of different resting patches visited per individual divided by the total amount of available patches *M* = 33 (a), and the fraction that represents the most visited patch (*M*_1_) over the whole amount of visited patches per individual (b). Higher values in saturation indicate higher exploration rates (more visited cluster over the period of study) while a larger fraction in *M*_1_ indicates a larger return rate to the most visited patch. These are snapshots at the end-time series for the LMEM used models that explain the exploration rate and the *M*_1_ distribution based on the variables: log-normalised time, age and sociality.

The saturation (equivalent to VUP as it describes the visited resting sites with respect to the maximum available patches) Figure 4 (a) could be statistically fit with a Linear Mixed-Effect Model Regression (LMEM), where we considered that some individuals could have repeated signals across multiple years. We also took into account how these data points grew in time and analysed their relationship with age and social information. The model showed how the triple interaction between log normalised time, age and sociality is a significant interaction in de-termining the observed pattern (*p-value <* 0.01). This reproduced a situation where sociality accelerates the exploration of different sites with time, but it is negatively coupled to age and therefore older and more social individuals exhibit a higher spatial fidelity, over younger cows which exhibit higher exploration rates.

In addition, the most visited cluster fraction Figure 4 was also fitted to a similar model (LMEM). Here we studied the cumulative occupancy of the most visited cluster (the curve *M*_1_ in Figure 4) and found similar results to the saturation analysis. In particular, every both social-ity and age were statistically significant (*p-value <* 0.01), with the interesting characteristic that even if sociality and age de-accelerate this grow in time (with a negative term), their force is mitigated by the triple interaction again, which again reproduces the situation where older and more social cows tend to have higher site fidelity and larger fractions of the most visited cluster.

### B. Model performance: reproducing empirical results

In order to understand how social effects can possibly drive the behaviour observed for resting choices, we tested our model in section III with respect to the data, and specifically we explored how the different parameters/details of the model should be tuned to reproduce the empirical patterns observed. For this, we run simulations using *M* = 30 and *N* = 60 (so reproducing approximately the empirical conditions discussed above).

#### 1. Frequency of imitation largely constrains the time scaling and the occupation distribution of visited patches

We first explored how the main properties of the occupation patterns change as a function of the imitation probability *p*. Figure 5 shows (from left to right) the results for (i) the average number of sites explored by each individual as a function of time, (ii) the correlation between pairs of individuals, defined as the histogram for the fraction of nights that two individuals share the same resting patch (averaged over all possible pairs of individuals), and (iii) the frequency of individuals sharing the same patch (averaged over all possible patches).

**FIG 5.**
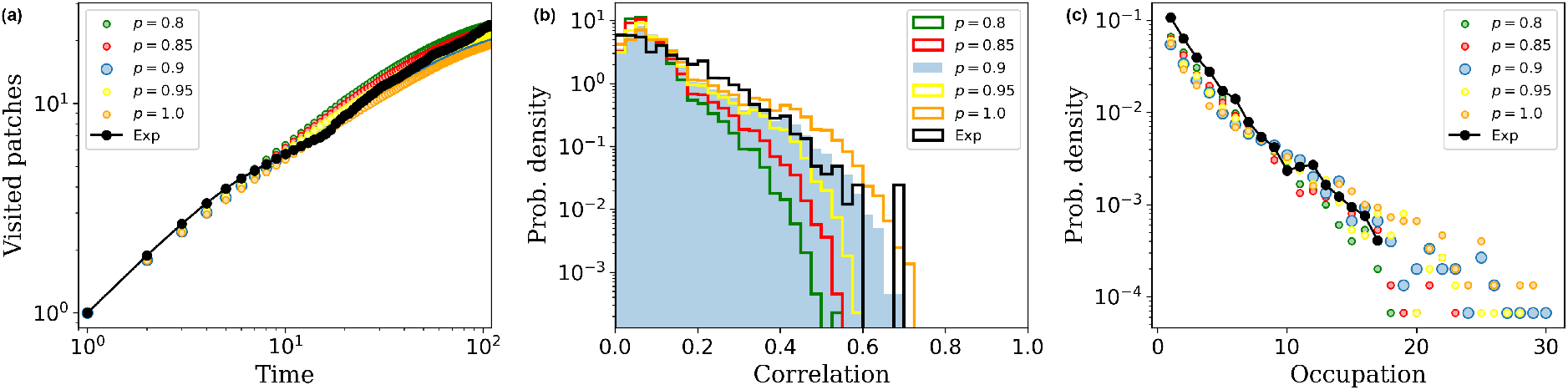
Exploration and occupation patterns obtained computationally from the model as a function of the imitation probability *p* (see legend) compared to the empirical data (exp, in black). The case *p* = 0.95 is stressed in the plots as it corresponds to the best fit obtained. The rest of the parameters used are *σ*_*p*_ = 0.4, *d* = 3, *N* = 60 and *M* = 30.

Our model suggests that high levels of imitation are required to fit the empirical data. As far as *p* takes values around 0.8 or higher, the exploration patterns (average number of different sites explored) as a function of time resemble those from the empirical data, and do not differ substantially when *p* is modified (Figure 5, left). Inter-estingly, the time dependence predicted by the model approximates to a power-law growth ⟨*s*⟩ ∼ *t*^0.6^, this is, the sub-linear regime observed in Figure 3, for a relatively wide range of the model parameters. We note, however, that this scaling is just a transient behaviour that emerges as a combination of initial exploration patterns driven by local imitation, and a latter stage of saturation where individuals have explored a significant set of patches, and the fraction of those still unexplored becomes gradually smaller (a direct comparison to the theoretical result in Eq. (12), not included here for simplicity, actually confirms this is the case).

Small imitation probabilities *p*, on the other side, yield very poor exploration (so the growth of new visited sites is very slow). Importantly, when *p* is not large enough we essentially have that random choices tend to dominate the dynamics of resting sites, in accordance to the discussion in section III A. As a result, the possibility that several cows share the same resting site becomes more and more unlikely. This is in contradiction to empirical data, which shows that (i) sites with an occupation of 10-15 individuals are relatively frequent (Figure 5, right) and (ii) correlations between individuals (measured as the fraction of nights that two specific individuals share their resting site) can take rather high values, even up to 0.4-0.5 (Figure 5, centre). At practice, we find that high values *p* ≈ 0.95 are necessary to fit the distribution of correlations and occupation patterns, which reduces random choices to only 5% of the cases.

#### 2. Random choices and individual variability are necessary to explain enhanced exploration patterns

In the preceding section, we have discussed why the average probability *p* must take large values. However, values of *p* too close to 1 are also detrimental, as imitation will be then the only mechanism driving site choices. So that, all individuals will tend to a ‘frozen’ situation with all individuals either sharing the same site or a few of them (separated by a distance larger than *d*, so local imitation cannot act between shared sites). In that case, abnormally large probabilities for high occupation appear. So, to avoid that situation we need that random choices, while being not dominant, have still a significant effect in the system.

Additionally, we find that when all individuals have the same value *p*_*i*_, then it is impossible to fit jointly the empirical exploration and occupancy patterns (this is, all empirical data in Figure 5). This means we need some levels of individual variability to explain such patterns. In particular, the best fits are obtained for a standard deviation *σ*_*p*_ = 0.4 combined with the mean *p* = 0.95 value above. The effect of shifting the value of *σ*_*p*_ above or below that optimal value can be observed in the sensitivity analysis of Figure 6. There we see that small values of this parameter yield slow exploration patterns (Figure 6, left) and too high occupation/sharing probabilities (Figure 6, centre and right). Instead, when the distribution of imitation probabilities is too broad (this is, for large *σ*_*p*_) then again random choices become dominant, and so high occupation probabilities rapidly decay to zero.

**FIG 6.**
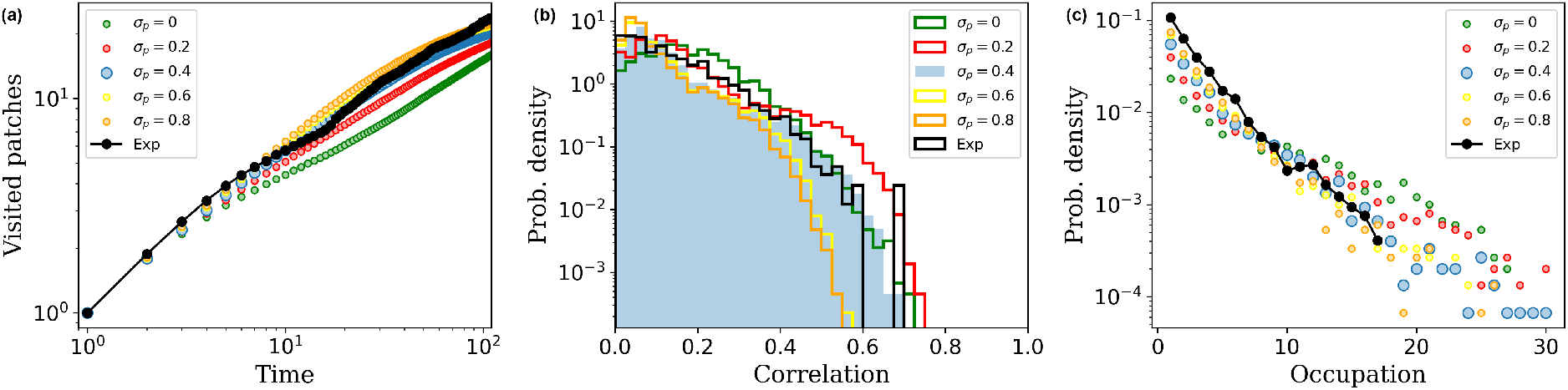
Exploration and occupation patterns obtained computationally from the model as a function of the dispersion *σ*_*p*_ in the imitation probability (see legend) compared to the empirical data (exp, in black). The case *σ*_*p*_ = 0.4 is stressed in the plots as it corresponds to the best fit obtained. The rest of the parameters used are *p* = 0.95, *d* = 3, *N* = 60 and *M* = 30.

#### 3. Local imitation is required to explain clustering effects in occupation

Finally, we stress that the fits presented in Figure 5 and Figure 6 correspond to the case *d* = 3, which is a relatively restrictive distance for imitation (note that *d* = 3 means that only six sites can be reached from the current one, while the system used has *M* = 30 available). Again, smaller or larger values of *d* rapidly lead to detrimental situations, as we show in Figure 7. For *d <* 3 we obtain again slow exploration patterns and too high probabilities for larger occupations, as it becomes difficult to the individuals to depart much from their current site. In the case *d* ≫ 3 we get the opposite effect: exploration becomes faster than that observed empirically, and the possibility that many individuals share the same site becomes extremely unlikely as now the mobility from any site to another becomes easier.

**FIG 7.**
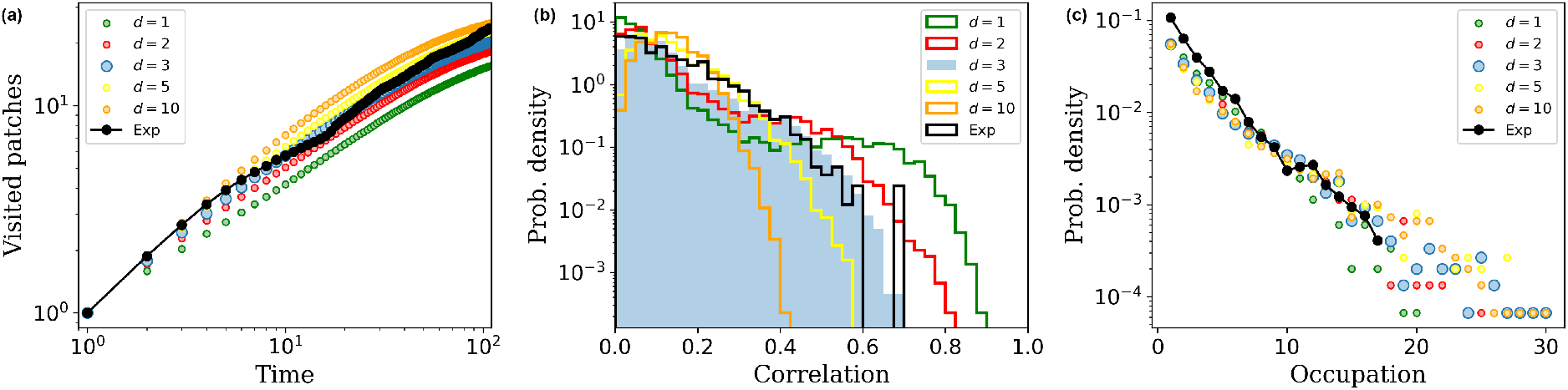
Exploration and occupation patterns obtained computationally from the model as a function of the maximum distance for imitation *d* (see legend) compared to the empirical data (exp, in black). The case *d* = 3 is stressed in the plots as it corresponds to the best fit obtained. The rest of the parameters used are *p* = 0.95, *σ*_*p*_ = 0.4, *N* = 60 and *M* = 30.

As a whole, this tells us that imitation effects must be essentially local in order to explain the empirical site choices, and we obtain a characteristic distance *d* = 3 for the effects of imitation. Restricting imitation locally makes sense if such imitation is to be interpreted as a mechanism for a gregarious behaviour, rather than one based on free-range individual prospection/copying, so enhancing the social nature of the dynamics observed.

#### 4. Revisit dynamics of cows reflects local imitation combined with the existence of preferred patches

Once we have checked the conditions that are required for the model to accommodate the empirical data, it is also convenient to discuss an additional feature we have obviated for the moment, as is the revisit dynamics of the individuals. As seen in Figure 3 and Figure 4, revisits to preferential sites are quite common in the dynamics of cows, reflecting a high persistence in the choices of resting sites. The most plausible interpretation for such high persistence is that it might be due to a combined effect of (i) memory effects (so individuals remember previously visited sites and have a tendency to revisit them), and/or (ii) variability between sites (this is, the existence of high-quality patches that are recurrently chosen for this reason). Our model in its present form is not able to explain such high levels of revisits, as it does not include any of these two effects. For this reason, we have explored alternative rules/mechanisms that may be included in the model for that purpose.

Essentially, we have explored the effect of introducing additional rules where individuals have a certain tendency to revisit some preferential places. To choose these preferential places, we have compared two situations: (a) preferential sites are different (and chosen at random) for each individual, or (b) preferential sites are the same for all individuals.

Remarkably, we have checked that rules of the type (a) are not able to explain empirical data. While it can increase the average number of revisits, including different preferential sites for every individual reduces significantly the correlations between individuals and reduce very much the probability of high occupancies. As a result, such models are not able to fit all our empirical results (exploration, occupation and revisit patterns) jointly.

Instead, we have checked that rules of type (b) add significant revisits (matching the empirical patterns observed) without distorting much the exploration and occupation dynamics. Altogether, this shows that heterogeneity between sites is another necessary ingredient to explain the dynamics of this social system.

We note that the heterogeneity between sites could be attributed to intrinsic differences between patches (high-quality vs low-quality patches), but it is also possible that they are due to some social factor (some patches may be more regularly used by individuals due to some conditioning, and for this reason they recurrently return to them). This confirms that social memory (rather than individual memory/experience) is the main driving force behind the revisit dynamics of the cows empirically observed.

## V. DISCUSSION

Resting patch selection is of critical importance in the steep, rugged study valley, as it can reduce mortality risk from falling on steep slopes and thus directly affect fitness [39]. Our findings show that cattle indeed select specific regions that are topographically flatter [30], in accordance with other studies in similar landscapes [40]. This selection can trigger a site fidelity behaviour, understood as re-visits to specific patches, which is evolutionary selected because of a lower uncertainty [5, 6, 24, 41], compared to dispersal where information has to be gathered from scratch [42, 43]. This landscape driven site fidelity behaviours, have already been observed in various environments [16, 44], although usually associated to highly predictable landscapes [45, 46]. In unpredictable environments, such as the study valley, where storms and sudden temperature and wind changes occur, but most resources are of high quality on average (such as water sources and grazing patches [30]), site fidelity behaviours have as well been observed [23, 45, 47].

Based on the precedent considerations, individual differences seem to play a significant role in decision-making [45, 48–5 Our findings show that individual hetero-geneities (measured as age and aspects of social behaviour), although statistically significant, have a small effect on the movement patterns across resting patches. Our data suggests that older and more social cows tend to have a lower exploration rate and a higher re-visit rate, which could indicate that their private information has a larger weight and that individual experience modulates site selection in the group [48, 50]. Therefore, the longer an individual is familiar with its habitat, the more prone it is to better select those resources across time and exhibits a stronger site fidelity behaviour [25, 50]. This experience, measured in terms of age, has profound fitness consequences [52, 53] and although most age-dependent responses are related to reproductive success [52, 54] or foraging [55, 56], our results hint at the idea that experience could improve the resting patch selection competence across individuals. However, these individual heterogeneities cannot entirely explain the observed collective patterns [8, 57, 58] and a different approach has to be taken.

Previous studies have shown that collective movement patterns can emerge from the interplay between individual differences and group-level decision rules [11, 19, 25,26, 49]. Our findings indicate that the exploration rate of resting patches grows in the sub-linear regime, with cattle visiting new sites at a smaller rate compared to that of random sampling. This pattern reflects a shift in the exploration–exploitation trade-off towards increased exploitation as long as exploration opportunities reduce or, alternatively, such exploration has already produced somewhat satisfactory choices. This hints at a higher expected revisit rate in cattle, as previously observed [24]. This behaviour has been usually associated to memory [19, 23, 25] or environmental factors [16, 59], but seldom to the social structure and dynamics of the group.

Consequently, we assumed here that for highly social species, the resting patch selection and fidelity was driven by a social copying mechanism instead of memory. Indeed, our social model quantitatively describes most of the observed patterns, based with just two simple rules: an imitation rule given by the social copying mechanism, and a inter-individual variability exploration given by a random sampling of the available patches. It suggests that sub-linear exploration dynamics can emerge from a social mechanism [4, 60]. Indeed, we observe how the sub-linearity and the cluster occupation distributions can be fairly matched when an imitation rule is introduced and it takes high imitation probabilities *p*. On top of that, inter-individual correlations are well explained and fit with empirical data, with some individuals showing high fidelity between them. This further suggests that simple social rules, such as social copying, can have a high impact on the decision making [5, 11, 61] and that public information plays a substantial role at population-level where the behaviour is amplified by the group’s social structure [3, 4, 6]. However, these have to be coupled to inter-individual heterogeneities (for example caused by age, experience, sociality or maternal status) to fully explain the observed behaviours [48, 51].

Interestingly, our model as it stands in Section III failed to predict the high site fidelity observed in our data. As discussed above, this could be caused by the additional effect of resting patch heterogeneities, as for example, distance to water sources or the grazing quality around resting patches, since previous studies have shown a relation between these heterogeneities and cattle behaviour [40, 62] . Such spatial heterogeneity between different resting patches could also cause the higher revisit rates [6, 16, 25, 47] and with direct fitness conse-quences [6].

Given the simplicity of the model, one could imagine that a feedback mechanism, coupling the resting patch selection with the social structure in the group could be observed in this case [4, 11]. Even if we did not directly explore this option, it is clear that by looking at the correlation between individuals, both in the empirical data and the ones predicted by the model that there is high fraction of individuals that share a significant amount of nights together Figure 5. This could potentially indicate that a higher social preference between individuals determines their respective movement decisions [49] and at the same time, the sharing of the same patch reinforces the social interaction, as an example of the feedback loop between the public and private information [4]. For these exposed reasons, our data suggest that individual differences can modulate the observed dynamical patterns [4, 48, 49, 51] but rely on the social copying mechanism to generate them [3, 11, 61]. This strengthens the idea that group characteristics and public information are more relevant in decision-making for highly social species [4, 6], with re-visits being explained through the social structure rather than individual memory.

## VI. CONCLUSIONS

Optimisation in resource selection through movement patterns is a critical step for improving fitness and survival aspects. Our findings strengthen the idea that complex dynamical patterns in resource selection might arise from the underlying social structure and a simple social copying process, which can be modulated by inter individual differences. Our study may open new avenues for the study of collective use of resources. An important question for future research is to determine under which ecological conditions social copying alone can generate recursive patterns of resource use, and when additional mechanisms such as individual memory become necessary to explain the observed collective dynamics. Our results also suggest the hypothesis that collective resource allocation may be disproportionately influenced by behavioural heterogeneity among individuals, even when average behavioural differences are weak. Testing how informed, experienced, or socially high-ranked individuals shape collective outcomes remains a challenge in studies of collective decision-making [63]. More generally, future studies should explore how social information, individual patterns, and environmental heterogeneity interact across spatio-temporal scales to generate the movement patterns commonly observed in collective systems. Understanding when these mechanisms reinforce or compensate one another may help fill the gap between current theoretical models and empirical observations [64–

## Data Availability

The datasets analysed during the current study are available in the *Dataset for: Social copying drives spatial allocation in gregarious herbivores and code* repository with DOI: 10.5281/zenodo.20646900.

## Acknowledgements

Funding was provided by the Next Generation UE program through the Spanish Ministry of Science (MCIU/AEI/FEDER, UE) (ref. TED2021-129425B-I00). DC acknowledges the financial support of the Spanish government through grant PID2021-122893NB-C22.

